# Identifying gastric cancer molecular subtypes by integrating DNA-based hierarchical classification strategy and clinical stratification

**DOI:** 10.1101/2023.06.09.544302

**Authors:** Binyu Yang, Siying Liu, Jiemin Xie, Xi Tang, Pan Guan, Yifan Zhu, Li C. Xia

## Abstract

**Background:** Molecular subtyping has been introduced to better understand the genetic landscape of gastric cancer (**GC**), but current subtyping methods only had limited success because of the mixed use of molecular features, a lack of strategy optimization, and the limited availability of GC samples. The community urgently calls for a precise, and easily adoptable subtyping method to enable DNA-based early screening and treatment.

**Methods:** Based on TCGA subtypes, we developed a novel classifier, termed Hierarchical DNA-based Classifier for Gastric Cancer Molecular Subtyping (**HCG**), leveraging all DNA-level alterations as predictors, including gene mutations, copy number aberrations and methylations. By adding the closely related esophageal adenocarcinomas (**EA**) dataset, we expanded the TCGA GC dataset for training and testing HCG (n=453). We optimized HCG with three hierarchical strategies evaluated by their overall accuracy using Lasso-Logistic regression, and by their clinical stratification capacity using multivariate survival analysis. We used difference tests to identify subtype-specific DNA alteration biomarkers based on HCG defined subtypes.

**Results:** Our HCG classifier achieved an overall AUC score of 0.95 and significantly improved the clinical stratification of patients (overall p-value=0.032). 25 subtype-specific DNA alterations were identified by difference tests, including high level of mutations in *SYNE1*, *ITGB4* and *COL22A1* genes for the MSI subtype, high level of methylations of *ALS2CL*, *KIAA0406* and *RPRD1B* genes for the EBV subtype.

**Conclusions:** HCG is an accurate and robust classifier for DNA-based GC molecular subtyping with high-performing clinical stratification capacity. The training and test datasets and analysis programs of HCG are available at https://github.com/labxscut/HCG.

## 1 INTRODUCTION

Gastric cancer (**GC**) is a globally important disease with over one million estimated new cases annually, ranking fifth in incidence rate and third in mortality rate worldwide [1]. Accurate subtyping of GC is essential for accurate diagnosis and prognosis, tumor staging, treatment guidance, recurrence monitoring, and drug development, as different subtypes exhibit distinct clinical outcomes and therapeutic responses [2,3]. Despite its importance, GC subtyping faces significant challenges due to the disease’s complex etiology and high molecular heterogeneity [2,4]. Therefore, developing effective GC subtyping methods is urgent and critical for improving the management and prognosis of patients with GC.

Before molecular subtyping, classification of GC was mainly based on microscopic histology features. One of the most widely used method is the Borrmann classification, which divides GC into four distinct subtypes (polypoid, fungating, ulcerated, and diffusely infiltrative carcinoma) just based on their gross appearance [5]. Another common method is the Lauren classification, which distinguishes between two histological subtypes (intestinal or diffusive) based on their microscopic features [6]. However, these non-molecular classifications have significant limitations in clinical practice, as the histological appearance may not adequately capture the biological diversity and heterogeneity of the underlying intrinsic mechanisms, so that these subtypes cannot provide meaningful basis for personalized therapy and precision medicine [7–9].

In order to reveal the underlying biological and molecular mechanisms of GC, researchers began to define more precise, effective and assessable GC subtypes at the molecular level [10,11]. The most impactful work was The Cancer Genome Atlas (**TCGA**)’s study, which classified GC into four subtypes: chromosomal instability (**CIN**), genomically stable (**GS**), microsatellite instability (**MSI**), and Epstein–Barr virus positive (**EBV**) [11], based on the genetic, epigenetic, and gene expression profiles of 295 primary GC cases. It was considered a seminal research to offer new perspectives for therapeutic development, and led to numerous follow-up works [12–14]. However, no consensus subtyping has yet been established.

The challenge to develop an accurate, robust and easily adoptable GC molecular subtyping classifier resides in several aspects. First, the use of molecular features in current multi-omics subtyping studies was undifferentiated and without biological rationale, leading to confusion in data acquisition, integration, and analysis [15–17]. Studies typically used genomics and transcriptomic features in a mixed manner. For instance, Cristescu *et al.* classified GC by integrating genomic and transcriptomic data [18]. Liu *et al.* applied a residual graph convolutional network to classify GC based on multi-omics fusion data [19]. Notably, the TCGA subtyping study also suffered from this issue. All these studies failed to consider the natural cascade of genetic information from genome to transcriptome and to proteome with increasing noise perturbation at lower levels [20]. Thus, mixed use of multi-level features adds unnecessary uncertainty to the resulting classifiers.

Second, there is no rigorous optimization of classification strategy by the studies so far. Primarily two types of subtyping strategies were used: one-step multi-class classification and multi-step hierarchical classification. One-step multi-class subtyping provides a simple and intuitive way to identify GC subtypes. For example, Zhou *et al.* simultaneously divided GCs into three subtypes in one step using 14 cancer functional states [21]. Li *et al.* clustered GCs into three subtypes (immunity-deprived (ImD), stroma-enriched (StE), and immunity-enriched (ImE)) based on the activities of 15 pathways associated with immune, DNA repair, oncogenic, and stromal signatures [3]. However, all these strategies did not consider the natural phylogeny of GC subtypes as a result of cancer development and progression, and risk not fully exploring this information for correct subtyping.

Hierarchical classification strategy was also frequently used. For example, The TCGA study categorized GC sequentially first by the presence of EBV features (EBV subtype), then by the presence of MSI signatures (MSI subtype), and finally by the number of somatic copy number alterations to genomically stable (GS subtype) or chromosomal instability (CIN subtype) subtypes [11]. For example, Tahara *et al.* classified GC into four subtypes, first by the CpG island methylator phenotype and then by the presence of TP53 hot spot mutations [22]. However, these hierarchical strategies varied arbitrarily in structure, subgrouping, and splitting. Moreover, all these multi-class and hierarchical strategies were adopted subjectively without scientific benchmark and justification. Therefore, a comprehensive and systematic way to evaluate these competing strategies is urgently needed.

Third, there were only limited public GC datasets available for developing a comprehensive multi-feature subtype classifier. In single-omics subtyping studies of GC, the most extensive methylation study is by Zhang *et al.* using 398 GC samples [23], and the most extensive transcriptome study is by Cristescu *et al.*, which assessed 300 GC samples’ gene expression data [24]. The largest multi-omics public data is from TCGA GC cohort, which had 383 samples with subtype information [25]. Therefore, there is significant need for more comprehensive datasets to improve our understanding of GC, and we would propose to aggregate samples from related cancers such as esophageal adenocarcinomas (**EA**), which most often occurs in the lower portion of the esophagus [26] and shares similar molecular subtypes with GC, to expand the training and testing sample availability.

Thus, we proposed a new approach to build a DNA alteration based GC subtype classifier, utilizing an expanded dataset and by systematically evaluating candidate classification strategies (**Figure 1A**). Our work is unique to develop a comprehensive DNA-level classifier. Compared to mRNA and protein changes, DNA alterations are more basic robust, and measurable. The central dogma dictates that genetic information flows from DNA to RNA and to protein [27], indicating that, for cancer cells, GC subtypes based on DNA features would be more fundamental. Moreover, DNA alterations are more reliable to ascertain, as DNA molecules are structurally stable and less susceptible to transcriptional and translational noises [28–31]. However, previous DNA-based subtyping studies did not use the full DNA-level information. For instance, Li *et al.* integrated only somatic mutational profiles and clinicopathological information to classify GC patients [32]; Usui G *et al.* used only DNA methylation patterns to classify GCs into distinct molecular subtypes [33]. Their partial use of DNA alterations may have neglected key features in left-out alteration types. Therefore, it is worthwhile to explore whether GC subtypes can be more accurately and robustly classified based on all available DNA alteration classes, including gene mutation, copy number aberration (**CNA**), and methylation alterations.

**Figure 1.**
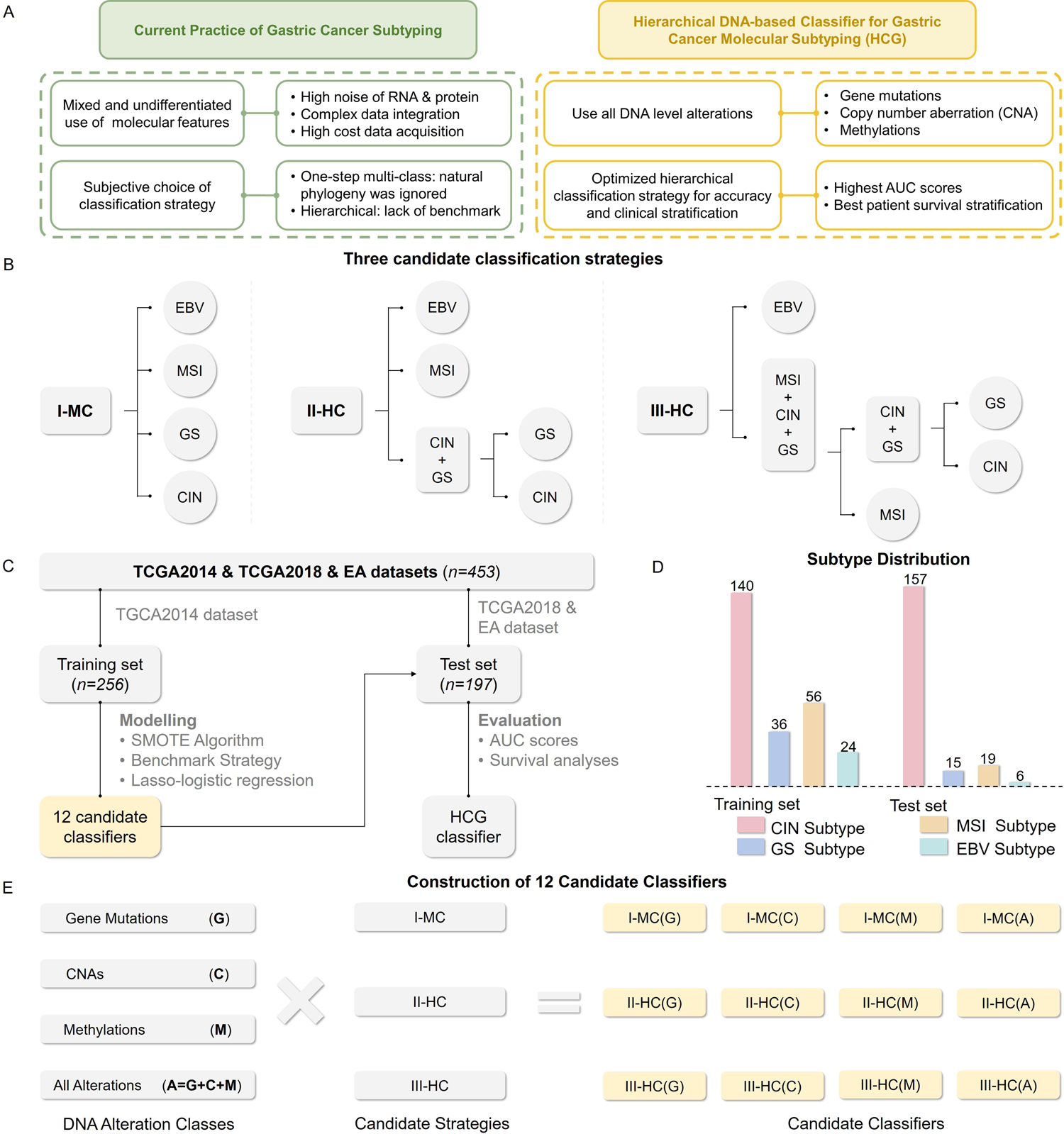
Study design, classification strategies and dataset description. **(A)** The conceptual design of DNA level GC subtyping. (A) Three candidate classification strategies evaluated in this study. **(C)** Training and validating process (TCGA2014 and TCGA2018 are both GC cohort). **(D)** Subtype distribution. **(E)** Enumeration of candidate classifiers.

Our work also aims to establish the criteria for evaluating the clinical efficacy and validity of competing molecular subtyping strategies. To ensure objectivity and eliminate potential biases in strategy selection, we conducted a comprehensive comparison of three candidate strategies (**Figure 1B**). We measured the classification accuracy using precision-recall curves. Besides, we evaluated the candidate classifiers’ clinical relevance, as this is crucial for their successful implementation in clinics [34].

Finally, our work aims to enlarge the sample space for multi-omics based GC subtype model. We integrated the TCGA EA dataset with the TCGA GC dataset to expand the sample number available for model training and testing. Our approach was supported by previous research highlighting significant similarities between EA and GC subtypes. Specifically, both diseases originate from the epithelial cells lining the digestive tract and they share similar risk factors and symptoms [35]. Additionally, molecular features such as chromosomal instability and TP53 mutations have been observed in both EA and GC [36]. Furthermore, immunotherapy efficacy is similar for EA and GC, primarily for those expressing PD-L1 or showing microsatellite instability [37]. Taking the CIN subtype as an example, which has the largest sample size in the EA dataset, we conducted a multivariate analysis of variance (MANOVA) test (see **Materials and Methods**), which showed no statistically significant difference between the CIN subtype of GC and EA datasets (p-value = 0.9462).

Our development accumulated to a novel DNA-level GC molecular subtype classifier **HCG** (**H**ierarchical DNA-based **C**lassifier for **G**astric Cancer Molecular Subtyping) (**Figure 1C**). It was developed on 453 combined GC+EA samples’ full DNA alterations (gene mutations, CNAs, and methylations) from the TCGA2014, TCGA 2018 and EA datasets, by rigorously benchmarking three classification strategies using Lasso-Logistic regression and multivariate survival analysis, and by adopting the best two-step hierarchical strategy selected. HCG demonstrated strong precision-recall performance (overall AUC=0.95) and the highest clinical stratification capacity (overall p-value=0.032). Based on the HCG subtypes, we performed difference tests and identified 25 statistically significant subtype-specific DNA alterations.

## 2 RESULTS

### 2.1 DNA alterations alone sufficiently and accurately identify gastric cancer subtypes

The total samples (n=453) analyzed in our study were first split into non-overlapping training (TCGA2014) and test (TCGA2018+EA) sets, as shown in **Figure 1D**. All the 12 DNA-level classifiers (**Figure 1E**) were trained and cross-validated using the training set augmented by the SMOTE (Synthetic Minority Oversampling Technique) algorithm [38], and then applied to the TCGA2018+EA test set. ROC (Receiver Operating Characteristic) curves and AUC (Area Under Curve) scores were used as classification performance evaluation metrics. Please refer to the **Materials and Methods** section for details.

We employed DNA-level mutation, CNA, methylation and their combination (all) as alteration classes to identify GC subtypes and evaluated their effectiveness using three candidate strategies (**Figure 1E**). The combination of input features and hierarchical learning strategies yielded 12 candidate classifiers, which we denoted as *S*_*i*_(*a*_*j*_), where *S*_*i*_ indicated the *i*-th classification strategy (*i* ∈{**I-MC**, **II-HC**, **III-HC**}, are abbreviations of One-step Multi-class Classification, Two-step Hierarchical Classification and Three-step Hierarchical Classification respectively) and *a*_*j*_ represented *j*-th input feature (*j* ∈{**G**, **C**, **M**, **A**}, which are abbreviations of gene mutations (G), CNAs (C), methylations (M) and all alteration classes (A) respectively). For instance, I-MC(G) denoted the classifier using gene mutations with the one-step multi-class (I-MC) strategy; II-HC(A) denoted the classifier using all classes of DNA alterations with the two-step hierarchical classification (II-HC) strategy.

We found that the classifiers using all alteration classes were more predictive than the corresponding ones using a single class on the test set. For instance, using the I-MC strategy, the overall AUC score for I-MC(A) was 0.95, indicating a 0.02, 0.05, and 0.03 increase over I-MC(G), I-MC(C), and I-MC(M) classifiers. We also observed similar trends for classifiers using the other strategies: Using the II-HC strategy, the overall AUC score for II-HC(A) was 0.95 (**Figure 2D**), indicating a 0.03, 0.10, and 0.04 increase over II-HC(G), II-HC(C) and II-HC(M) classifiers, respectively; using the III-HC strategy, the overall AUC score for III-HC(A) was 0.96, indicating a 0.03, 0.08, and 0.04 increase over III-HC(G), III-HC(C), and III-HC(M) classifiers, respectively. Consequently, the use of all alteration classes was significantly critical, enabling highly accurate identification of GC subtypes with overall AUC >0.95 in all subtypes.

**Figure 2.**
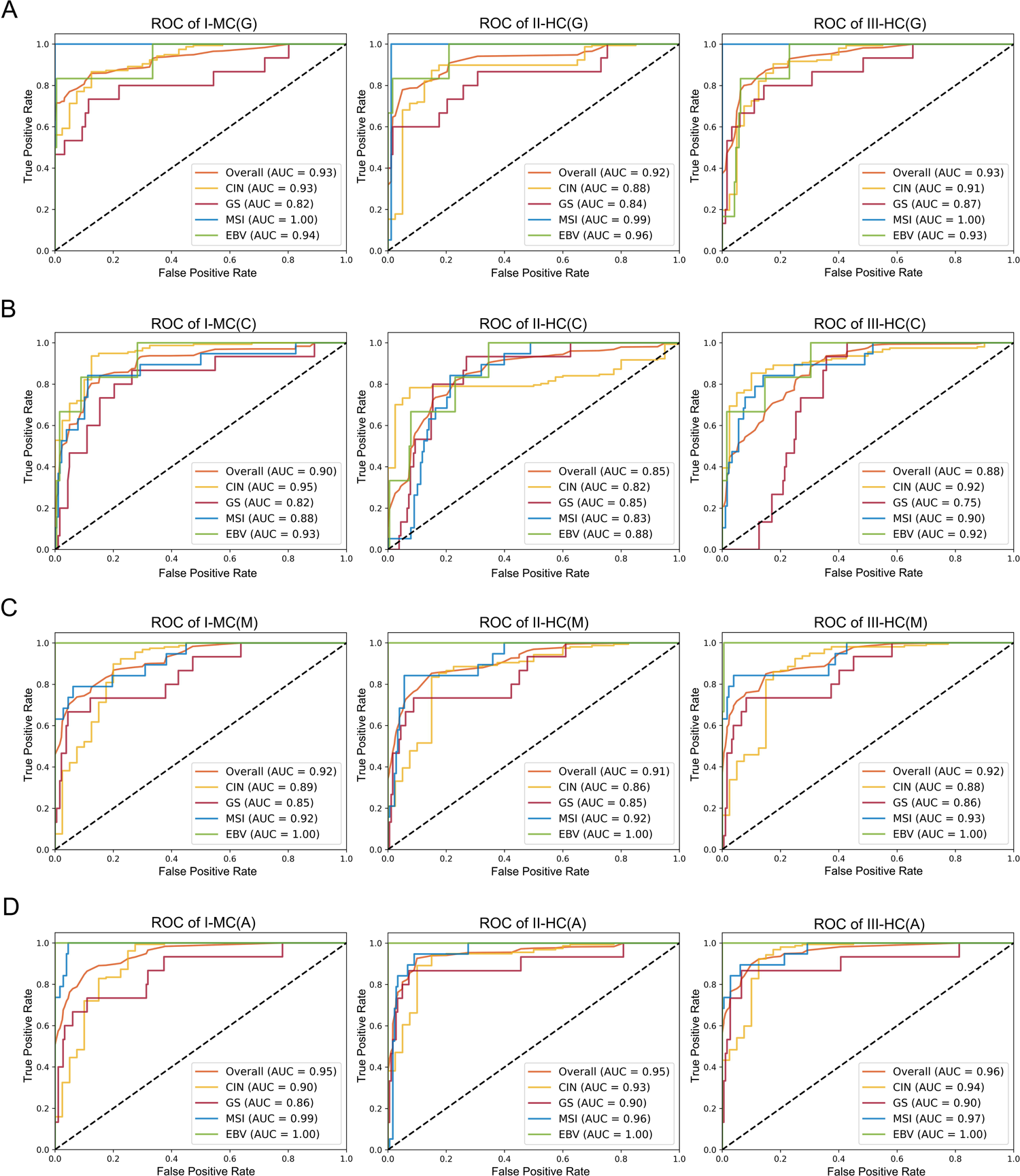
ROC curves of 12 candidate classifiers on test set. **(A)** ROC curves for Ⅰ-MC(G), Ⅱ-HC(G), Ⅲ-HC(G). **(B)** ROC curves for Ⅰ-MC (C), Ⅱ-HC(C), Ⅲ-HC(C). **(C)** ROC curves for Ⅰ-MC(M), Ⅱ-HC(M), Ⅲ-HC(M). **(D)** ROC curves for Ⅰ-MC(A), Ⅱ-HC(A), Ⅲ-HC(A).

We also found that one class of DNA alterations can already accurately predict subtype in a few cases. For example, I-MC(G), II-HC(G), and III-HC(G) strategies using gene mutations only performed the best in predicting the MSI subtype, with almost perfect AUC scores 1.00, 0.99 and 1.00, respectively (**Figure 2A**). While the AUC scores of gene-mutation only classifiers were significantly lower, with 0.93, 0.88 and 0.91 respectively for the CIN subtype, 0.82, 0.84 and 0.87 for the GS subtype, 0.94, 0.96 and 0.93 for the EBV subtype. This suggested that point mutations are the main alterations driving the development of the MSI subtype but not other subtypes, which aligns with the nature of the MSI subtype-arising from DNA mismatch repair (MMR) gene malfunctions and being characterized by extremely high level of point mutations [39].

Similarly, I-MC(M), II-HC(M), and III-HC(M) strategies using methylations only performed the best in predicting the EBV subtype, which all had perfect AUC score 1.00 (**Figure 2C**). In contract, methylation-based classifiers had significantly lower AUC scores of 0.89, 0.86 and 0.88 respectively for the CIN subtype, 0.85, 0.85 and 0.86 respectively for the GS subtype, and 0.92, 0.92 and 0.93 respectively for the MSI subtype. This indicated a substantial difference between EBV methylation profiles and the other subtypes; As an external validation, previous studies reported abnormally higher levels of methylation in EBV subtype, suggesting that methylation features are of vital importance in defining EBV subtype of GC as we observed [40].

### 2.2 Two-step hierarchical classifier II-HC(A) demonstrates the best clinical stratification

In light of the superior performance using all classes of DNA alterations, we conducted a comparative analysis evaluating candidate classification strategies configured in this way. Overall, the I-MC(A), II-HC(A) and III-HC(A) classifiers all achieved highly competitive overall AUC scores (0.95, 0.95, and 0.96, respectively). This trend was similar for subtype-specific AUC score (see **Figure 2**). Specifically, these classifiers all obtained the perfect AUC scores of 1.00 for the EBV subtype, 0.90, 0.93, and 0.94 for the CIN subtype; 0.86, 0.90, and 0.90 for the GS subtype; and 0.99, 0.96, and 0.97 for the MSI subtype, respectively (**Figure 2D**). Notably, all these classifiers achieved the highest accuracy for the EBV subtype, followed by the MSI, and then CIN and GS subtypes, which suggested CIN and GS subtypes are the most difficult subtypes to resolve.

To evaluate their clinical relevance, we performed multivariate survival analyses on subtypes predicted by the original TCGA study, and I-MC(A), II-HC(A), III-HC(A) classifiers (see **Materials and Methods**). The analyses included the patient age and sex. We set the reference as subtype=CIN, age<65 and sex=female. The results were presented in **Table 1**, while those concerning age and sex were provided in **Supplementary Table 1**. Our analyses revealed that the II-HC(A) classifier demonstrated the best clinical stratification capacity (overall p-value=0.032 for the multivariate Cox model, and a reduction of 0.019, 0.014, and 0.025 in comparison with TCGA, I-MC(A), and III-HC(A) defined subtypes, respectively.). Specifically, the II-HC(A) classifier achieved the most statistically significant stratification for patients with the EBV subtype (p-value=0.221) and the second most statistically significant stratification among patients with the GS and MSI subtypes (p-value=0.551 and 0.064, respectively). This is generally better than the TCGA results—the current golden standard. Considering their overall accuracy and clinical relevance, we adopted II-HC(A) as the underlying classifier for HCG.

### 2.3 HCG subtypes are linked to specific molecular features in gastric cancer

We found that HCG subtypes were substantially different from TCGA defined subtypes. The confusion matrix and the alluvial plot (**Figure 3A, B**) revealed that subtype-switching occurred in 9.71% (44 out of 453) of the samples analyzed. Here we denoted the HCG defined subtype X as HCG-X and TCGA defined subtype Y as TCGA-Y. We observed that the TCGA-EBV subtype remained unchanged upon HCG classification. However, the TCGA-GS subtype demonstrated the most significant transition, with 47.06% of it switching to HCG-CIN, 5.88% to HCG-MSI, and 1.96% to HCG-EBV. Furthermore, a small fraction of the TCGA-CIN subtype was found to be HCG-GS (0.34%) and HCG-MSI (2.02%). Similarly, 12% TCGA-MSI subtype was found switching to HCG-CIN (10.67%) and HCG-EBV (1.33%).

**Figure 3.**
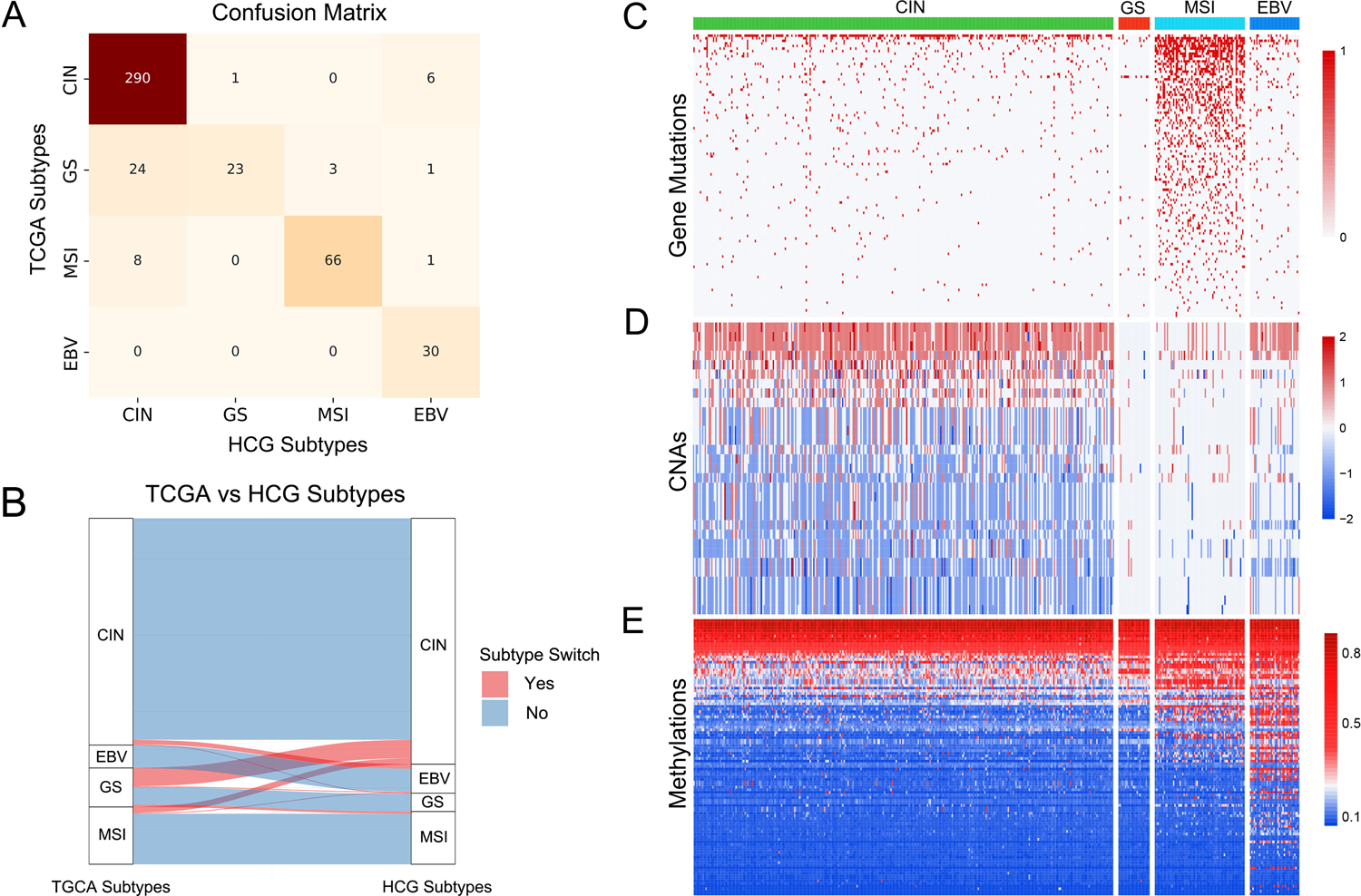
Analysis of HCG subtypes. **(A)** Confusion matrix between TCGA subtypes and HCG subtypes. **(B)** Alluvial plot between TCGA subtypes and HCG subtypes. **(C-E)** Heatmaps of all significant gene mutations, CNAs, and methylations.

We found that almost 50% of TCGA-GS subtype switched to HCG-CIN subtype, which may explain HCG’s significant improvement is clinical stratification. Previous studies have highlighted that CIN tumor cells inhibit the immune response and exhibit greater chemotherapy sensitivity as compared to GS tumor cells, leading to significant differences in treatment response and patient prognosis [3,13]. This may lead to substantial survival differences between the two GC subtypes. Therefore, the more accurate identification of CIN and GS molecular subtypes using the HCG classifier may have significant implication in clinics.

To visualize the genome-wide DNA alteration patterns by subtypes, we generated a heatmap based on 247 DNA alteration features as selected by HCG (**Figure 3C–3E**). We observed that the HCG-MSI subtype indeed exhibited the highest frequency of mutations (**Figure 3C**). Specifically, the mutation rates (see **Materials and Methods**) for the HCG-CIN, GS, MSI, and EBV subtypes were found to be 0.0123, <u>0.0044</u>,0.0998, and 0.0140, respectively, where its highly elevated in MSI. This result corroborated with numerous previous researches suggesting that the emergence of MSI subtype is linked to dysfunctional DNA mismatch repair pathway, often leading to the occurrence of large amounts of point mutations [40].

Moreover, we found that the HCG-CIN subtype had the highest CNA rate (see **Materials and Methods**), while the HCG-GS subtype had the lowest (**Figure 3D**). The computed CNA rates for HCG-CIN, GS, MSI, and EBV subtypes were 0.544, <u>0.0245</u>,0.097, and 0.0433, respectively, which aligned with numerous previous research that identified the CIN subtype as associated with extensive chromosomal loss or rearrangement, whereas the GS subtype was characterized as being genomically stable (i.e. absent of large-scale CNAs) [41–43].

Furthermore, we found that the HCG-EBV subtype displayed hypermethylation characteristics (**Figure 3E**). The methylation rates (see **Materials and Methods**) for the HCG-CIN, GS, MSI, and EBV subtypes were <u>0.254</u>, 0.271, 0.281, and 0.326, respectively. Notably, only the HCG-EBV subtype exhibited elevated hypermethylation pattern, which greatly surpassed the average methylation rate 0.266. The result also aligned with earlier studies highlighting the extreme DNA hypermethylation that characterizes the EBV subtype [33,40,44].

### 2.4 The HCG classifier identifies novel subtype-specific DNA alterations

Discovering subtype-specific markers is crucial for developing targeted therapies. Taking advantage of HCG’s better classification results, we performed difference tests (see **Materials and Methods**) and identified 25 statistically significant subtype-specific DNA alterations, as shown in **Table 2**.

For the HCG-CIN subtype, 3 CNA-based (*CDH1*, *CDH3*, *KIF26B*) and 2 methylation-based (*CDR2*, *CUL4B*) DNA alteration markers were identified. Indeed, there are reports associating these DNA alterations with GC in general. For example, *CDH1* gene encodes E-cadherin and its mutations, are the most common germline mutations detected in GC, underlying hereditary diffuse gastric cancer [45]. *CDH3* encodes P-cadherin and its up-regulation was associated with GC [46]. *KIF26B* encodes a kinesin family member and is one of the most upregulated genes in metastatic GC samples [47]. *CUL4B* promotes cell invasion and epithelial–mesenchymal transition in vitro, as well as tumor growth and metastasis in vivo [48]. But our analysis was among the first to validate their CIN-specific nature. Notably, the HCG classifier also newly identified *CDR2* hypomethylation as CIN-specific marker. Previously, *CDR2* was identified as a tumor antigen expressed in a high percentage of breast cancer [49]. However, the mechanism behind *CDR2* and CIN subtype association remains to be explored.

For the HCG-GS subtype, 10 mutation-based (*NFASC*, *MYOM2*, *PTPRJ*, *PLXNB2*, *TTK*, *ECEL1*, *ARHGEF15*, *TRPM4*, *CYP4F2*, *CA10*) and 4 methylation-based (*PRKCQ*, *IGF1*, *CRYGB*, *TSTD1*) subtype markers were identified. Notably, the *TSTD1* hypermethylation was newly identified by the HCG classifier as the GS-specific marker. According to previous studies, *TSTD1* expression was significantly negatively correlated with *FZD1*, while *FZD1* showed a significant positive correlation with *FZD2* and *FZD6*, which were all associated with GC. In fact, high expression of *FZDs* could inhibit the proliferation, migration and invasion of GC cells by activating the non-classical Wnt signaling pathway [50]. Therefore, the hypermethylation of *TSTD1* may reduce its expression, thus promote the expression of *FZD* genes, which deserves further validation.

For the HCG-MSI subtype, our analysis identified 3 signature alterations, including high level point mutation in the *SYNE1*, *ITGB4*, and *COL22A1* genes. These are consistent with the increased mutational load inherent to the MSI subtype [51]. Notably, the mutated *ITGB4* gene was newly identified by the HCG classifier as an MSI-specific marker. In fact, the MSI subtype is driven by underlying defects in the DNA mismatch repair system [52] which results in various dysfunctions in the Wnt pathway, including the Wnt/β-catenin signaling pathway, that affects cell proliferation [53]. It has been shown that presence of *ITGB4* mutations decrease or impair the encoded integrin β4 subunit [54]. The β4 subunit participants in the Wnt/β-catenin signaling pathway by inhibiting Wnt-responsive gene transcription and reducing cell division [55]. Therefore, the mutated *ITGB4* gene may cause its low expression or dysfunctions, thus lose its repressive effect on the Wnt signaling pathway [52,56], which in turn enhances the MSI phenotype.

For the HCG-EBV subtype, our analysis identified 3 signature alterations, including *RPRD1B*, *KIAA0406* and *ALS2CL* hypermethylation. These are consistent with high CpG island methylator phenotype of the EBV subtype [57]. The product of *RPRD1B* gene was reported to accelerate the G2/M transition by interacting with Aurora kinase B and the ROS-related p53 pathway, thereby facilitating faster division of GC cells [58]. Therefore, the hypermethylation of *RPRD1B* can suppress cancer cell division, which makes it a protective factor for EBV subtypes. This observation is consistent with the better prognosis associated with the EBV subtype [13]. As another example, the *KIAA0406* gene encodes a vital constituent of the TTT complex, known to regulate the DNA damage response [59]. Hypermethylation of *KIAA0406* may lead to decreases the functionality of the TTT complex, permitting increased DNA damage and mutational burdens that aligns with the observed higher mutational burdens also observed in the EBV subtype (second highest mutation rate, next to MSI) [57].

## 3 DISCUSSION

Molecular subtypes and signatures were associated with distinct clinical outcomes in general. They are being delineated in various solid tumors, thereby laying the groundwork for improved clinical management via precision-driven treatment plans [60–62]. For gastric cancer, several molecular subtyping studies have been reported [10,13,32,44,63]. Yet no consensus has been reached, which calls for novel model development. Here we developed an accurate, robust, and easily adoptable molecular subtype classifier HCG, which demonstrated high subtyping accuracy and significantly improved clinical stratification of GC patients.

Our approach represented a more robust and reproducible way to identify subtypes in GC. Previous studies have shown that integrating multi-omics data to identify subtypes can make relatively accurate prediction, such as TCGA [11] and many others [18–19]. Nevertheless, with this study we demonstrated that DNA-level information is the most significant and sufficient source in GC subtype classification. The AUC scores for the HCG classifier using all DNA alterations categories were all greater than 0.95. There are biological reasons to our observation, as DNA-level genetics and epigenetics changes determine downstream phenotypes. For example, DNA methylation intrinsically modulates chromatin structure and has a protein-independent regulatory role that increases the stiffness of DNA, determines the 3D structure of the genome, and as a result regulates the gene expression [64–65]. Therefore, aberrations in the genome and epigenome are expected to synergistically interact and contribute to gastric carcinogenesis, while further research and comprehensive analyses combing both are needed to fully understand the extent of this deterministic relationship.

Our study also pointed out that the consideration of clinical relevance is critical in evaluating subtyping classifiers. The classification structure profoundly influences the clinical relevance in general. For example, Pretzsch *et al.* established a classification system for molecular subtyping of GC using immunohistochemistry and morphology-based analyses as markers, which had clinical relevance and reproduced the ACRG molecular subtypes of GC [66]. Ramos *et al.* found that IHC/ISH analysis was able to distinguish immunophenotypic groups of GC with distinct characteristics and prognosis [67]. In our study, we also found varying clinical relevance among the different competitively performing classifiers. Specifically, the survival outcomes were better stratified based on the HCG (II-HC(A)) subtypes (overall p-value = 0.032) compared to TCGA subtypes (overall p-value = 0.051), while the stratification based on III-HC(A) subtypes (overall p-value = 0.057) were inferior to those of TCGA, though the II-HC(A) and III-HC(A) classifiers had similar accuracy. Besides, correct clinical stratification can provide crucial information that influences therapeutic decisions and patient outcomes [68,69]. Therefore, it is vital to assess classifiers with regards to clinical relevance.

Our study supported that molecular subtypes should be considered when developing new GC drugs and therapies, as they all demonstrated significant subtype-specific DNA-level alterations. Indeed, clinical trials have reported that MSI-H status may be a biomarker for pembrolizumab therapy among patients. Specifically, among patients with the MSI subtype of GC, the median overall survival (OS) was not reached for both pembrolizumab monotherapy (95% CI, 10.7 months to not reached) and pembrolizumab plus chemotherapy (95% CI, 3.6 months to not reached) compared with a median OS of 8.5 months (95% CI, 5.3-20.8 months) for chemotherapy alone. The early divergence of the survival curves between patients with the MSI subtype who received pembrolizumab therapy vs chemotherapy suggests that earlier introduction of pembrolizumab is beneficial in patients with the MSI subtype of GC [70]. As another example, several studies suggested that chemotherapy might not be the optimal treatment choice for patients with the MSI subtype. Previous studies have demonstrated that patients with MSI subtype respond worse to chemotherapy alone [71]. Adverse reactions have also been observed in patients with the MSI subtype who underwent neoadjuvant chemotherapy. Even, patients with the MSI subtype who responded to neoadjuvant chemotherapy did not experience enhanced prognoses compared to non-responders [72]. Therefore, with improved patient subtyping, HCG has the potential to guide and enhance GC treatment.

Finally, our study saw limitations. First, the public data available for our analysis was limited. Despite our attempts to address the inadequacy of available data by additionally incorporating the EA samples, the need for an augmented volume of data persists. Furthermore, we encountered an issue with sample imbalance as the majority of the EA samples were of the CIN subtype. While this was mitigated by SMOTE balancing in training, further targeted studies in minor subtypes are required. Second, since the HCG classifier was designed relying on the four GC molecular subtypes defined by the TCGA project, other subtype classification schemes may be not yet viable in our model. In fact, researchers have reported several GC molecular subtype classification methods [3,10], which could be considered in our future models as their data become more available.

## 4 MATERIALS AND METHODS

### 4.1 Data source and pre-processing

The genomic data analyzed in this study, including gene mutations, copy number alterations (CNAs), methylation alterations and clinical data, were obtained from the cBioPortal database (https://www.cbioportal.org/) [73]. The sample labels of CIN, GS, MSI and EBV molecular subtypes were derived from the TCGA Gastric Cancer (GC) study in 2014 and denoted as TCGA2014. Furthermore, an additional dataset from the TCGA PanCanAtlas published in 2018 was included and denoted as TCGA 2018. The TCGA dataset including esophageal adenocarcinomas (EA) samples were denoted as EA (**Figure 1C**).

TCGA2014 was used as the training set, while TCGA2018 and EA were used as test sets. To pre-process the mutation data, the downloaded mutation profiles were mapped to the gene level using a Boolean matrix where each cell indicated whether the *i* − *th* gene was mutated at least once in the *j* − *th* patient. For CNAs, copy number values downloaded from cBioPortal were mapped to the gene level. For methylation data, missing values were filled with the nearest ten neighbors by applying the *knnImputation()* function in R’s *DMwR* package if there were no more than half missing [74–76]. Genes corresponding to multiple methylation values were then assigned the averaged value. Finally, we standardized each alteration using Min-Max Scaling, as shown in the following formula:

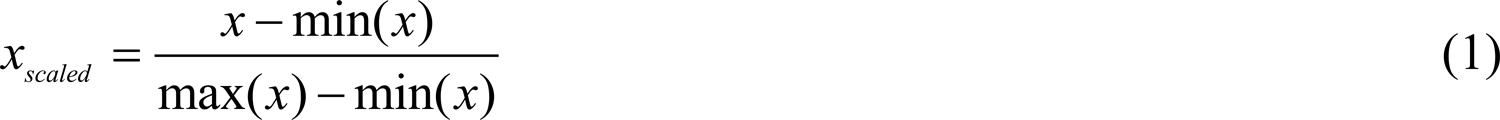

where *x* represents the value of each alteration, while min(*x*) and max(*x*) denote the minimum and maximum values of the alteration across all samples, respectively.

### 4.2 SMOTE algorithm

To address the issue of imbalanced distribution in the four subtypes (shown in **Figure 1D**) and to mitigate potential biases, we applied a data augmentation technique called Synthetic Minority Oversampling Technique (**SMOTE**) prior to training [38]. Specifically, we increased the number of samples for the GS, MSI and EBV subtypes in the training set by generating synthetic samples using the k-nearest neighbor method. This approach resulted in an increase number of training samples (balanced for 140 samples for each subtype) and reduced the risk of overfitting in the constructed model. The SMOTE algorithm was implemented using the *imblearn* library in Python.

### 4.3 Lasso-Logistic regression

We applied the Lasso-Logistic regression model to solve the classification problem, consisting of Least Absolute Shrinkage and Selection Operator (Lasso) and the logistic regression methods, which is implemented in Python through the scikit-learn library. Lasso is an efficient approach to model high-dimensional and multi-modal data, and it avoids overfitting by soft thresholding, which selects significant alteration features by regularizing the regression coefficients of non-informative features to zeros. While implementing the model, we performed a 5-fold cross-validation. For the regularization weight *λ* (lambda), we chose the minimal lambda that achieves the highest classification prediction accuracy.

In details, suppose the response variable has *K* levels *G* = {1, 2,…, *K*}, and each sample *x_i_* in our study has *m* (*m* = 53496) features, i.e., *x_i_ = (x_i,1_, x_i,2_, …, x_i,m_)*, then we model:

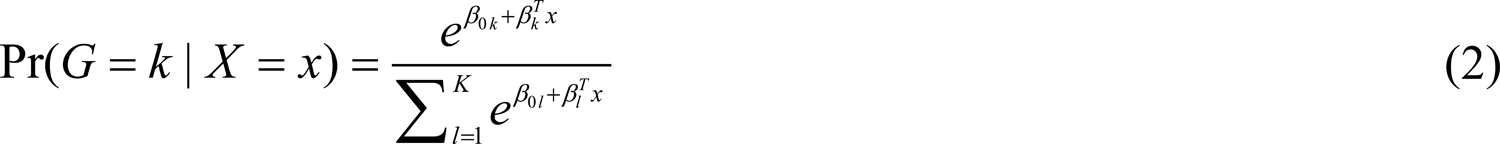

Let *Y* be the *N* × *K* indicator response matrix, with elements *y_il_* = *I* (*g_i_* = *l*). Then the Lasso elastic net penalized negative log-likelihood function of logistic regression becomes:

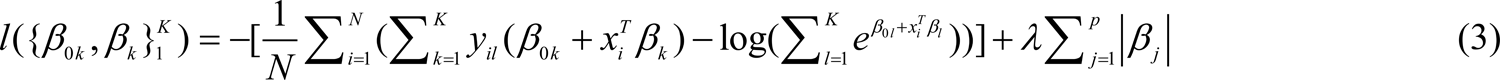

where *β* is a *p* × *K* matrix of coefficients, *_k_* refers to the *k* − *th* column (for outcome category *k*), and the *j* − *th* row (vector of *K* coefficients for variable *j*), *λ* is the penalty parameter as chosen by previous cross-validation.

### 4.4 ROC curve and AUC score

The performances of the 12 candidate classifiers were assessed by Receiver Operating Characteristic (ROC) curves and the Area Under the Curve (AUC) scores. The ROC curve is a graphical tool used to display the performance of a classifier. It shows the sensitivity and specificity of a classifier at different thresholds. The AUC score, in turn, can be used as a numerical measure to intuitively evaluate the general performance of a classifier, with larger values indicating better performance.

While the ROC curve is primarily used to evaluate binary classification, it can be extended to multi-class classification using the One-vs-Rest (**OVR**) strategy. This approach involves treating each class as the positive class while all other classes are considered the negative. By employing this OVR method, we generated ROC curve for each class, and the classifier’s ability to distinguish a particular class can then be assessed by its corresponding AUC score. The overall performance of the classifier was evaluated by calculating the average of the AUC scores for each class.

### 4.5 Survival analysis

To evaluate the clinical stratification ability of the classifiers, survival analyses were performed in R using the *survival* and *survminer* package with Overall Survival (**OS**) as the primary endpoint. In univariate analysis, the Log-rank test was used to compare survival outcomes and Kaplan-Meier (**KM**) curves were generated to provide a visual representation of the survival data.

In multivariate analysis, patient age and sex were included as a covariate. Cox proportional hazards regression was used to analyze the effect of subtypes, age and sex, using the *coxph()* R function. The forest plot (organized into **Table 1** and **Supplementary Table 1**) showed survival outcomes with subtype=CIN, age<65 and sex=male as reference. Analyses involving multiple comparisons were adjusted using the Benjamini-Hochberg (BH) FDR method, which controls the False Discovery Rate (FDR) with the Holm–Bonferroni method for multiple hypothesis testing. All statistical tests were determined to be significant based on the BH-corrected significance level of 0.05.

### 4.6 Mutation, CNA and methylation rates

For mutation rate, we first computed the average mutation rate of each sample across all genes. Then, we used the HCG classifier to categorize the samples into *k* subtype (*k* ∈{*CIN*,*GS*, *MSI*, *EBV*}) and calculated the average mutation rate of each subtype. Let

*mu_i_*, *_j_* be the mutation count for sample *i* in gene *j* and be the total number of genes in mutation profiles. The mutation rate for the *k* − *th* subtype is:

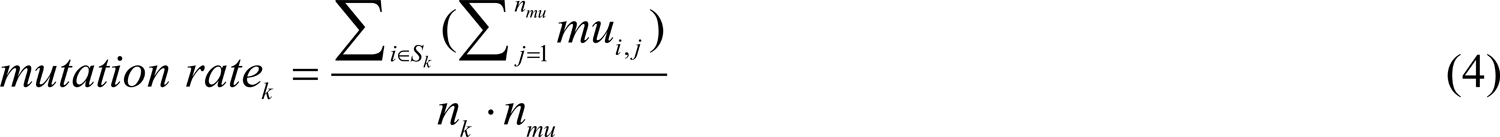

where *S* is the set of the *k* − *th* subtype and *n* is the number of samples belong to the *k* − *th* subtype.

Similarly, we defined the CNA and methylation rates in Equations (6) and (7). Let *c_i_* _, *j*_ and *me_i_*, *_j_* be the copy number aberration count and methylation level for sample *i* in gene *j*, respectively, ^n^CNA and ^n^me be the total number of genes in CNA and methylation profiles, respectively. Then the CNA rate and methylation rate for the *k* − *th* subtype are:

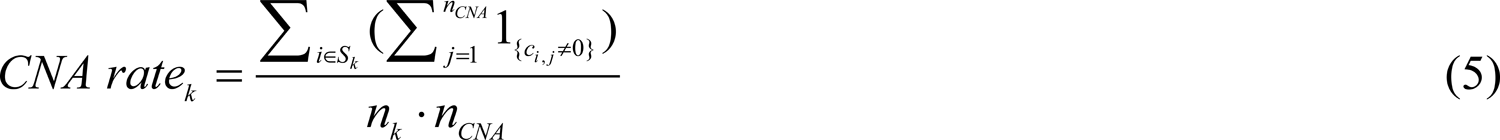

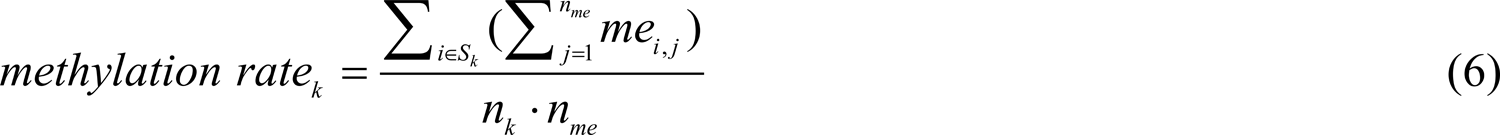

where 1(·) is indicator function.

### 4.7 Difference test

In order to identify significant subtype-specific DNA alterations, difference tests were conducted in R on 247 DNA alterations selected by the HCG classifier. For mutation data, the Fisher’s exact test was applied using the *fisher.test()* R function. For copy number aberrations and methylation data, the independent Student’s T-test was employed using the *t.test*() R function. DNA alterations were considered significantly different if the adjusted p-value (FDR correction method) was less than 0.00001 using a two-sided test with the OvR strategy. Finally, we performed duplicate gene removal.

To compare differences between the CIN subtype of GC and EA datasets, we conducted multivariate analysis of variance (MANOVA) using the *manova()* R function. MANOVA test allows for the simultaneous analysis of multiple dependent variables and is used to determine if there are any significant differences between groups based on multiple response variables.

## ACKNOWLEDGEMENTS

This study was funded by the Guangdong Basic and Applied Basic Research Foundation (2022A1515-011426 to L.C.X.), National Natural Science Foundation of China (61873027 to L.C.X.)

## COMPLIANCE WITH ETHICS GUIDELINES

The authors declare that they have no conflict of interest or financial conflicts to disclose. The article does not contain any human or animal subjects performed by any of the authors.

## AUTHOR CONTRIBUTIONS

B.Y., S.L., J.X., X.T. and L.C.X. conceived and designed the study. S.L. and Y.Z. collected the data. B.Y., S.L., and P.G. performed the modelling and analysis. B.Y., S.L., J.X. and L.C.X. wrote the manuscript. All authors edited the manuscript and approved the submitted version.

## ABBREVIATIONS

GC: Gastric cancer

EA: Esophageal adenocarcinomas

TCGA: The Cancer Genome Atlas

CIN: Chromosomal instability

GS: Genomically stable

MSI: Microsatellite instability

EBV: Epstein–Barr virus positive

CNA: Copy number aberration

HCG: Hierarchical DNA-based classifier for gastric cancer molecular subtyping

ROC: Receiver operating characteristic

AUC: Area under curve

OvR: One-vs-Rest

OS: Overall survival

I-MC: One-step multi-class classification strategy

II-HC: Two-step hierarchical classification strategy

III-HC: Three-step hierarchical classification strategy

I-MC(A): One-step multi-class classification strategy with all classes of alterations

I-MC(G): One-step multi-class classification strategy with gene mutations

I-MC(M): One-step multi-class classification strategy with methylations

II-I-MC(C): One-step multi-class classification strategy with CNAs

III-HC(A): Two-step hierarchical classification strategy with all classes of alterations

II-HC(G): Two-step hierarchical classification strategy with gene mutations

II-HC(M): Two-step hierarchical classification strategy with methylations

II-HC(C): Two-step hierarchical classification strategy with CNAs

III-HC(A): Three-step hierarchical classification strategy with all classes of alterations

III-HC(G): Three-step hierarchical classification strategy with gene mutations

III-HC(M): Three-step hierarchical classification strategy with methylations

III-HC(C): Three-step hierarchical classification strategy with CNAs

SMOTE: Synthetic minority oversampling technique

LASSO: Least absolute shrinkage and selection operator

MANOVA: Multivariate analysis of variance

---

**Supplementary Figure 1.** Alluvial plots. **(A)** Alluvial plot between TCGA subtypes and I-MC(A) subtypes. **(B)** Alluvial plot between TCGA subtypes and III-HC(A) subtypes.

**Table. 1** Multivariate survival analyses using all samples from TCGA and I-MC(A), II-HC(A) and III-HC(A) defined subtypes.

**Table. 2** Novel subtype-specific DNA alterations as identified by HCG.

**Supplementary Table. 1** Multivariate survival analyses using all samples from TCGA and I-MC(A), II-HC(A) and III-HC(A) defined subtypes (Age and Sex).

## Notes

### Competing Interest Statement

The authors have declared no competing interest.

## REFERENCES

1. Smyth, E. C., Nilsson, M., Grabsch, H. I., van Grieken, N. C., and Lordick, F. (2020) Gastric cancer. The Lancet. 396, 635–648

2. Gao, J.-P., Xu, W., Liu, W.-T., Yan, M., and Zhu, Z.-G. (2018) Tumor heterogeneity of gastric cancer: From the perspective of tumor-initiating cell. World Journal of Gastroenterology. 24, 2567

3. Li, L., and Wang, X. (2021) Identification of gastric cancer subtypes based on pathway clustering. npj Precision Oncology. 5, 46

4. Alsina, M., Gullo, I., and Carneiro, F. (2017) Intratumoral heterogeneity in gastric cancer: A new challenge to face. Annals of Oncology. 28, 912–913

5. Borrmann, R. (1926) Geschwülste des magens und duodenums. Verdauungsschlauch: Rachen and Tonsillen· Speiseröhre Magen and Darm· Bauchfell. 812–1054

6. Lauren, P. (1965) The two histological main types of gastric carcinoma: Diffuse and so-called intestinal-type carcinoma. An attempt at a histo-clinical classification. Acta Pathol Microbiol Scand. 64, 31–49

7. Garattini, S. K., Basile, D., Cattaneo, M., Fanotto, V., Ongaro, E., Bonotto, M., Negri, F. V., Berenato, R., Ermacora, P., Cardellino, G. G., et al. (2017) Molecular classifications of gastric cancers: Novel insights and possible future applications. World J Gastrointest Oncol. 9, 194–208

8. Serra, O., Galán, M., Ginesta, M. M., Calvo, M., Sala, N., and Salazar, R. (2019) Comparison and applicability of molecular classifications for gastric cancer. Cancer Treat Rev. 77, 29–34

9. Wang, Q., Liu, G., and Hu, C. (2019) Molecular classification of gastric adenocarcinoma. Gastroenterology Res. 12, 275–282

10. Lei, Z., Tan, I. B., Das, K., Deng, N., Zouridis, H., Pattison, S., Chua, C., Feng, Z., Guan, Y. K., and Ooi, C. H. (2013) Identification of molecular subtypes of gastric cancer with different responses to pi3-kinase inhibitors and 5-fluorouracil. Gastroenterology. 145, 554–565

11. Network, C. G. A. R. (2014) Comprehensive molecular characterization of gastric adenocarcinoma. Nature. 513, 202

12. Katona, B. W., and Rustgi, A. K. (2017) Gastric cancer genomics: Advances and future directions. Cellular and Molecular Gastroenterology and Hepatology. 3, 211–217

13. Sohn, B. H., Hwang, J.-E., Jang, H.-J., Lee, H.-S., Oh, S. C., Shim, J.-J., Lee, K.-W., Kim, E. H., Yim, S. Y., and Lee, S. H. (2017) Clinical significance of four molecular subtypes of gastric cancer identified by the cancer genome atlas project. Clinical Cancer Research. 23, 4441–4449

14. Nshizirungu, J. P., Bennis, S., Mellouki, I., Sekal, M., Benajah, D.-A., Lahmidani, N., El Bouhaddouti, H., Ibn Majdoub, K., Ibrahimi, S. A., and Celeiro, S. P. (2021) Reproduction of the cancer genome atlas (tcga) and asian cancer research group (acrg) gastric cancer molecular classifications and their association with clinicopathological characteristics and overall survival in moroccan patients. Disease Markers.

15. Wörheide, M. A., Krumsiek, J., Kastenmüller, G., and Arnold, M. (2021) Multi-omics integration in biomedical research–a metabolomics-centric review. Analytica Chimica Acta. 1141, 144–162

16. Pinu, F. R., Beale, D. J., Paten, A. M., Kouremenos, K., Swarup, S., Schirra, H. J., and Wishart, D. (2019) Systems biology and multi-omics integration: Viewpoints from the metabolomics research community. Metabolites. 9, 76

17. Voillet, V., Besse, P., Liaubet, L., San Cristobal, M., and González, I. (2016) Handling missing rows in multi-omics data integration: Multiple imputation in multiple factor analysis framework. BMC Bioinformatics. 17, 1–16

18. Röcken, C. (2017) Molecular classification of gastric cancer. Expert Rev Mol Diagn. 17, 293–301

19. Liu, C., Duan, Y., Zhou, Q., Wang, Y., Gao, Y., Kan, H., and Hu, J. (2022) A classification method of gastric cancer subtype based on residual graph convolution network. Frontiers in Genetics. 13,

20. Raser, J. M., and O’shea, E. K. (2005) Noise in gene expression: Origins, consequences, and control. Science. 309, 2010–2013

21. Zhou, Q., Yuan, Y., Lu, H., Li, X., Liu, Z., Gan, J., Yue, Z., Wu, J., Sheng, J., and Xin, L. (2023) Cancer functional states-based molecular subtypes of gastric cancer. Journal of Translational Medicine. 21, 80

22. Tahara, T., Tahara, S., Horiguchi, N., Okubo, M., Terada, T., Yamada, H., Yoshida, D., Omori, T., Osaki, H., Maeda, K., et al. (2019) Molecular subtyping of gastric cancer combining genetic and epigenetic anomalies provides distinct clinicopathological features and prognostic impacts. Hum Mutat. 40, 347–354

23. Lian, Q., Wang, B., Fan, L., Sun, J., Wang, G., and Zhang, J. (2020) DNA methylation data-based molecular subtype classification and prediction in patients with gastric cancer. Cancer Cell International. 20, 349

24. Cristescu, R., Lee, J., Nebozhyn, M., Kim, K.-M., Ting, J. C., Wong, S. S., Liu, J., Yue, Y. G., Wang, J., Yu, K., et al. (2015) Molecular analysis of gastric cancer identifies subtypes associated with distinct clinical outcomes. Nature Medicine. 21, 449–456

25. Sanchez-Vega, F., Mina, M., Armenia, J., Chatila, W. K., Luna, A., La, K. C., Dimitriadoy, S., Liu, D. L., Kantheti, H. S., and Saghafinia, S. (2018) Oncogenic signaling pathways in the cancer genome atlas. Cell. 173, 321–337. e310

26. Coleman, H. G., Xie, S.-H., and Lagergren, J. (2018) The epidemiology of esophageal adenocarcinoma. Gastroenterology. 154, 390–405

27. Crick, F. (1970) Central dogma of molecular biology. Nature. 227, 561–563

28. Watson, J. D., and Crick, F. H. (1953) Molecular structure of nucleic acids: A structure for deoxyribose nucleic acid. Nature. 171, 737–738

29. Franklin, R. E., and Gosling, R. G. (1953) Evidence for 2-chain helix in crystalline structure of sodium deoxyribonucleate. Nature. 172, 156–157

30. Breslauer, K. J., Frank, R., Blöcker, H., and Marky, L. A. (1986) Predicting DNA duplex stability from the base sequence. Proceedings of the National Academy of Sciences. 83, 3746–3750

31. Kool, E. T., Morales, J. C., and Guckian, K. M. (2000) Mimicking the structure and function of DNA: Insights into DNA stability and replication. Angewandte Chemie International Edition. 39, 990–1009

32. Li, X., Wu, W. K. K., Xing, R., Wong, S. H., Liu, Y., Fang, X., Zhang, Y., Wang, M., Wang, J., Li, L., et al. (2016) Distinct subtypes of gastric cancer defined by molecular characterization include novel mutational signatures with prognostic capability. Cancer Research. 76, 1724–1732

33. Usui, G., Matsusaka, K., Mano, Y., Urabe, M., Funata, S., Fukayama, M., Ushiku, T., and Kaneda, A. (2021) DNA methylation and genetic aberrations in gastric cancer. Digestion. 102, 25–32

34. Zhao, L., Lee, V. H., Ng, M. K., Yan, H., and Bijlsma, M. F. (2019) Molecular subtyping of cancer: Current status and moving toward clinical applications. Briefings in Bioinformatics. 20, 572–584

35. Hayakawa, Y., Sethi, N., Sepulveda, A. R., Bass, A. J., and Wang, T. C. (2016) Oesophageal adenocarcinoma and gastric cancer: Should we mind the gap? Nature Reviews Cancer. 16, 305–318

36. Salem, M. E., Puccini, A., Xiu, J., Raghavan, D., Lenz, H. J., Korn, W. M., Shields, A. F., Philip, P. A., Marshall, J. L., and Goldberg, R. M. (2018) Comparative molecular analyses of esophageal squamous cell carcinoma, esophageal adenocarcinoma, and gastric adenocarcinoma. The Oncologist. 23, 1319–1327

37. Shankaran, V., Xiao, H., Bertwistle, D., Zhang, Y., You, M., Abraham, P., and Chau, I. (2021) A comparison of real-world treatment patterns and clinical outcomes in patients receiving first-line therapy for unresectable advanced gastric or gastroesophageal junction cancer versus esophageal adenocarcinomas. Advances in Therapy. 38, 707–720

38. Chawla, N. V., Bowyer, K. W., Hall, L. O., and Kegelmeyer, W. P. (2002) Smote: Synthetic minority over-sampling technique. Journal of Artificial Intelligence Research. 16, 321–357

39. Ratti, M., Lampis, A., Hahne, J. C., Passalacqua, R., and Valeri, N. (2018) Microsatellite instability in gastric cancer: Molecular bases, clinical perspectives, and new treatment approaches. Cellular and Molecular Life Sciences. 75, 4151–4162

40. Wang, K., Yuen, S. T., Xu, J., Lee, S. P., Yan, H. H., Shi, S. T., Siu, H. C., Deng, S., Chu, K. M., and Law, S. (2014) Whole-genome sequencing and comprehensive molecular profiling identify new driver mutations in gastric cancer. Nature Genetics. 46, 573–582

41. Rajagopalan, H., Nowak, M. A., Vogelstein, B., and Lengauer, C. (2003) The significance of unstable chromosomes in colorectal cancer. Nature Reviews Cancer. 3, 695–701

42. Geigl, J. B., Obenauf, A. C., Schwarzbraun, T., and Speicher, M. R. (2008) Defining ‘chromosomal instability’. Trends in Genetics. 24, 64–69

43. Ling, Y., Watanabe, Y., Nagahashi, M., Shimada, Y., Ichikawa, H., Wakai, T., and Okuda, S. (2020) Genetic profiling for diffuse type and genomically stable subtypes in gastric cancer. Computational and Structural Biotechnology Journal. 18, 3301–3308

44. Jácome, A. A. d. A., Lima, E. M. d., Kazzi, A. I., Chaves, G. F., Mendonça, D. C. d., Maciel, M. M., and Santos, J. S. d. (2016) Epstein-barr virus-positive gastric cancer: A distinct molecular subtype of the disease? Revista da Sociedade Brasileira de Medicina Tropical. 49, 150–157

45. Luo, W., Fedda, F., Lynch, P., and Tan, D. (2018) Cdh1 gene and hereditary diffuse gastric cancer syndrome: Molecular and histological alterations and implications for diagnosis and treatment. Frontiers in Pharmacology. 9, 1421

46. Imai, K., Hirata, S., Irie, A., Senju, S., Ikuta, Y., Yokomine, K., Harao, M., Inoue, M., Tsunoda, T., and Nakatsuru, S. (2008) Identification of a novel tumor-associated antigen, cadherin 3/p-cadherin, as a possible target for immunotherapy of pancreatic, gastric, and colorectal cancers. Clinical Cancer Research. 14, 6487–6495

47. Zhang, H., Ma, R., Wang, X., Su, Z., Chen, X., Shi, D., Guo, X., Liu, H., and Gao, P. (2017) Kif26b, a novel oncogene, promotes proliferation and metastasis by activating the vegf pathway in gastric cancer. Oncogene. 36, 5609–5619

48. Qi, M., Jiao, M., Li, X., Hu, J., Wang, L., Zou, Y., Zhao, M., Zhang, R., Liu, H., and Mi, J. (2018) Cul4b promotes gastric cancer invasion and metastasis-involvement of upregulation of her2. Oncogene. 37, 1075–1085

49. O’Donovan, K. J., Diedler, J., Couture, G. C., Fak, J. J., and Darnell, R. B. (2010) The onconeural antigen cdr2 is a novel apc/c target that acts in mitosis to regulate c-myc target genes in mammalian tumor cells. PLoS One. 5, e10045

50. Li, Y., Liu, Z., and Zhang, Y. (2021) Expression and prognostic impact of fzds in pancreatic adenocarcinoma. BMC Gastroenterology. 21, 79

51. Di Bartolomeo, M., Morano, F., Raimondi, A., Miceli, R., Corallo, S., Tamborini, E., Perrone, F., Antista, M., Niger, M., and Pellegrinelli, A. (2020) Prognostic and predictive value of microsatellite instability, inflammatory reaction and pd-l1 in gastric cancer patients treated with either adjuvant 5-fu/lv or sequential folfiri followed by cisplatin and docetaxel: A translational analysis from the itaca-s trial. The Oncologist. 25, e460–e468

52. Hinata, M., and Ushiku, T. (2021) Detecting immunotherapy-sensitive subtype in gastric cancer using histologic image-based deep learning. Scientific Reports. 11, 22636

53. Nusse, R., and Clevers, H. (2017) Wnt/β-catenin signaling, disease, and emerging therapeutic modalities. Cell. 169, 985–999

54. Nakano, A., Pulkkinen, L., Murrell, D., Rico, J., Lucky, A. W., Garzon, M., Stevens, C. A., Robertson, S., Pfendner, E., and Uitto, J. (2001) Epidermolysis bullosa with congenital pyloric atresia: Novel mutations in the β4 integrin gene (itgb4) and genotype/phenotype correlations. Pediatric Research. 49, 618–626

55. Rima, M., Daghsni, M., Lopez, A., Fajloun, Z., Lefrancois, L., Dunach, M., Mori, Y., Merle, P., Brusés, J. L., and De Waard, M. (2017) Down-regulation of the wnt/β-catenin signaling pathway by cacnb4. Molecular Biology of The Cell. 28, 3699–3708

56. Wang, J., Li, R., He, Y., Yi, Y., Wu, H., and Liang, Z. (2020) Next-generation sequencing reveals heterogeneous genetic alterations in key signaling pathways of mismatch repair deficient colorectal carcinomas. Modern Pathology. 33, 2591–2601

57. Chen, Z.-H., Yan, S.-M., Chen, X.-X., Zhang, Q., Liu, S.-X., Liu, Y., Luo, Y.-L., Zhang, C., Xu, M., and Zhao, Y.-F. (2021) The genomic architecture of ebv and infected gastric tissue from precursor lesions to carcinoma. Genome Medicine. 13, 1–22

58. Jia, Y., Yan, Q., Zheng, Y., Li, L., Zhang, B., Chang, Z., Wang, Z., Tang, H., Qin, Y., and Guan, X.-Y. (2022) Long non-coding rna neat1 mediated rprd1b stability facilitates fatty acid metabolism and lymph node metastasis via c-jun/c-fos/srebp1 axis in gastric cancer. Journal of Experimental & Clinical Cancer Research. 41, 1–20

59. Hurov, K. E., Cotta-Ramusino, C., and Elledge, S. J. (2010) A genetic screen identifies the triple t complex required for DNA damage signaling and atm and atr stability. Genes & Development. 24, 1939–1950

60. Higgins, M. J., and Baselga, J. (2011) Targeted therapies for breast cancer. The Journal of Clinical Investigation. 121, 3797–3803

61. Roepman, P., Schlicker, A., Tabernero, J., Majewski, I., Tian, S., Moreno, V., Snel, M. H., Chresta, C. M., Rosenberg, R., and Nitsche, U. (2014) Colorectal cancer intrinsic subtypes predict chemotherapy benefit, deficient mismatch repair and epithelial-to-mesenchymal transition. International Journal of Cancer. 134, 552–562

62. Li, T., Kung, H.-J., Mack, P. C., and Gandara, D. R. (2013) Genotyping and genomic profiling of non–small-cell lung cancer: Implications for current and future therapies. Journal of Clinical Oncology. 31, 1039

63. Ahn, S., Lee, S.-J., Kim, Y., Kim, A., Shin, N., Choi, K. U., Lee, C.-H., Huh, G. Y., Kim, K.-M., and Setia, N. (2017) High-throughput protein and mrna expression–based classification of gastric cancers can identify clinically distinct subtypes, concordant with recent molecular classifications. The American Journal of Surgical Pathology. 41, 106–115

64. Bonev, B., and Cavalli, G. (2016) Organization and function of the 3d genome. Nature Reviews Genetics. 17, 661–678

65. Buitrago, D., Labrador, M., Arcon, J. P., Lema, R., Flores, O., Esteve-Codina, A., Blanc, J., Villegas, N., Bellido, D., Gut, M., et al. (2021) Impact of DNA methylation on 3d genome structure. Nat Commun. 12, 3243

66. Pretzsch, E., Bösch, F., Todorova, R., Nieß, H., Jacob, S., Guba, M., Kirchner, T., Werner, J., Klauschen, F., and Angele, M. K. (2022) Molecular subtyping of gastric cancer according to acrg using immunohistochemistry–correlation with clinical parameters. Pathology-Research and Practice. 231, 153797

67. Ramos, M. F. K. P., Pereira, M. A., de Mello, E. S., dos Santos Cirqueira, C., Zilberstein, B., Alves, V. A. F., Ribeiro-Junior, U., and Cecconello, I. (2021) Gastric cancer molecular classification based on immunohistochemistry and in situ hybridization: Analysis in western patients after curative-intent surgery. World Journal of Clinical Oncology. 12, 688

68. Tandirerung, F. J. (2022) The clinical importance of differentiating monogenic familial hypercholesterolemia from polygenic hypercholesterolemia. Current Cardiology Reports. 1–9

69. Khushman, M. d., Patel, G. K., Maharjan, A. S., McMillin, G. A., Nelson, C., Hosein, P., and Singh, A. P. (2021) The prevalence and clinical relevance of 2r/2r tyms genotype in patients with gastrointestinal malignancies treated with fluoropyrimidine-based chemotherapy regimens. The Pharmacogenomics Journal. 21, 308–317

70. Chao, J., Fuchs, C. S., Shitara, K., Tabernero, J., Muro, K., Van Cutsem, E., Bang, Y.-J., De Vita, F., Landers, G., and Yen, C.-J. (2021) Assessment of pembrolizumab therapy for the treatment of microsatellite instability–high gastric or gastroesophageal junction cancer among patients in the keynote-059, keynote-061, and keynote-062 clinical trials. JAMA Oncology. 7, 895–902

71. Kim, S. Y., Choi, Y. Y., An, J. Y., Shin, H. B., Jo, A., Choi, H., Seo, S. H., Bang, H. J., Cheong, J. H., and Hyung, W. J. (2015) The benefit of microsatellite instability is attenuated by chemotherapy in stage ii and stage iii gastric cancer: Results from a large cohort with subgroup analyses. International Journal of Cancer. 137, 819–825

72. Cai, Z., Rui, W., Li, S., Fingerhut, A., Sun, J., Ma, J., Zang, L., Zhu, Z., and Zheng, M. (2020) Microsatellite status affects tumor response and survival in patients undergoing neoadjuvant chemotherapy for clinical stage iii gastric cancer. Frontiers in Oncology. 10, 614785

73. Cerami, E., Gao, J., Dogrusoz, U., Gross, B. E., Sumer, S. O., Aksoy, B. A., Jacobsen, A., Byrne, C. J., Heuer, M. L., and Larsson, E. (2012) The cbio cancer genomics portal: An open platform for exploring multidimensional cancer genomics data. Cancer Discovery. 2, 401–404

74. Fleischer, T., Frigessi, A., Johnson, K. C., Edvardsen, H., Touleimat, N., Klajic, J., Riis, M. L., Haakensen, V. D., Wärnberg, F., and Naume, B. (2014) Genome-wide DNA methylation profiles in progression to in situand invasive carcinoma of the breast with impact on gene transcription and prognosis. Genome Biology. 15, 1–13

75. Suh, Y.-S., Na, D., Lee, J.-S., Chae, J., Kim, E., Jang, G., Lee, J., Min, J., Ock, C.-Y., and Kong, S.-H. (2022) Comprehensive molecular characterization of adenocarcinoma of the gastroesophageal junction between esophageal and gastric adenocarcinomas. Annals of Surgery. 275, 706–717

76. Yang, X., Gao, L., and Zhang, S. (2017) Comparative pan-cancer DNA methylation analysis reveals cancer common and specific patterns. Briefings in Bioinformatics. 18, 761–773

