## Supplementary material for "Identifying gastric cancer molecular subtypes by integrating DNA-based hierarchical classification strategy and clinical stratification": Table 1

| **Table. 1** Multivariate survival analyses using all samples from TCGA and I-MC(A), II-HC(A) and III-HC(A) defined subtypes. | | | | | | |
| --- | --- | --- | --- | --- | --- | --- |
| **Classifier** | **Subtype** | **Count (%)** | **95% CI** | **Hazard Ratio** | **P-value^1^** | **Overall Stratification P-value^2^** |
| TCGA | CIN (reference) | 285 (66.6%) | / | / | / | 0.051 |
|  | GS | 47 (11.0%) | (0.60, 1.63) | 0.99 | 0.973 |  |
|  | MSI | 66 (15.4%) | (0.41, 1.01) | 0.64 | 0.056 |  |
|  | EBV | 30 (7.0%) | (0.41, 1.51) | 0.79 | 0.468 |  |
| I-MC(A) | CIN (reference) | 316 (73.8%) | / | / | / | 0.046* |
|  | GS | 37 (8.6%) | (0.75, 2.04) | 1.23 | 0.412 |  |
|  | MSI | 58 (13.6%) | (0.39, 1.04) | 0.64 | 0.07 |  |
|  | EBV | 17 (4.0%) | (0.40, 2.07) | 0.91 | 0.825 |  |
| II-HC(A) | CIN (reference) | 309 (72.2%) | / | / | / | **0.032*** |
|  | GS | 22 (5.1%) | (0.63, 2.39) | 1.23 | 0.551 |  |
|  | MSI | 59 (13.8%) | (0.40, 1.01) | 0.64 | 0.064 |  |
|  | EBV | 38 (8.9%) | (0.38, 1.22) | 0.69 | 0.221 |  |
| III-HC(A) | CIN (reference) | 312 (72.9%) | / | / | / | 0.057 |
|  | GS | 23 (5.4%) | (0.64, 2.43) | 1.24 | 0.524 |  |
|  | MSI | 64 (15.0%) | (0.42, 1.10) | 0.67 | 0.088 |  |
|  | EBV | 29 (6.7%) | (0.42, 1.53) | 0.80 | 0.495 |  |
| ^1^Statistical significance is based on the fitted multivariate cox model (log-rank test).  ^2^Statistical significance is based on the fitted multivariate cox model (likelihood ratio test), * if P<0.05. | | | | | | |
