## Supplementary material for "Identifying gastric cancer molecular subtypes by integrating DNA-based hierarchical classification strategy and clinical stratification": Table 2

| **Table. 2** Novel subtype-specific DNA alterations as identified by HCG. | | | | |
| --- | --- | --- | --- | --- |
|  | **CIN** | **GS** | **MSI** | **EBV** |
| **Gene Mutation** | / | *NFASC*,  *MYOM2*,  *PTPRJ*,  *PLXNB2**,  *TTK*,  *ECEL1**, *ARHGEF15**,  *TRPM4**,  *CYP4F2**,  *CA10* | *SYNE1*, *ITGB4**,  *COL22A1** | / |
| **Copy number aberration** | *CDH1*(+), *CDH3*(+),  *KIF26B*(-) | / | / | / |
| **Methylation** | *CDR2*(-)*, *CUL4B*(+) | *PRKCQ*(-)*,  *IGF1*(-),  *CRYGB*(-)*,  *TSTD1*(+)* | / | *ALS2CL*(+)*, *KIAA0406*(+)*,  *RPRD1B*(+) |
| Note: markers in each grid were ordered by statistical significance as found by difference tests.  (+) denotes copy number amplification or hypermethylation. (-) denotes copy number deletion or hypomethylation.  *: were not previously associated with GC to our knowledge | | | | |
