## Supplementary material for "Identifying gastric cancer molecular subtypes by integrating DNA-based hierarchical classification strategy and clinical stratification": GC_supplementary file

**Supplementary Materials includes:**

1. Supplementary Figures

2. Supplementary Tables

**1. Supplementary Figures**

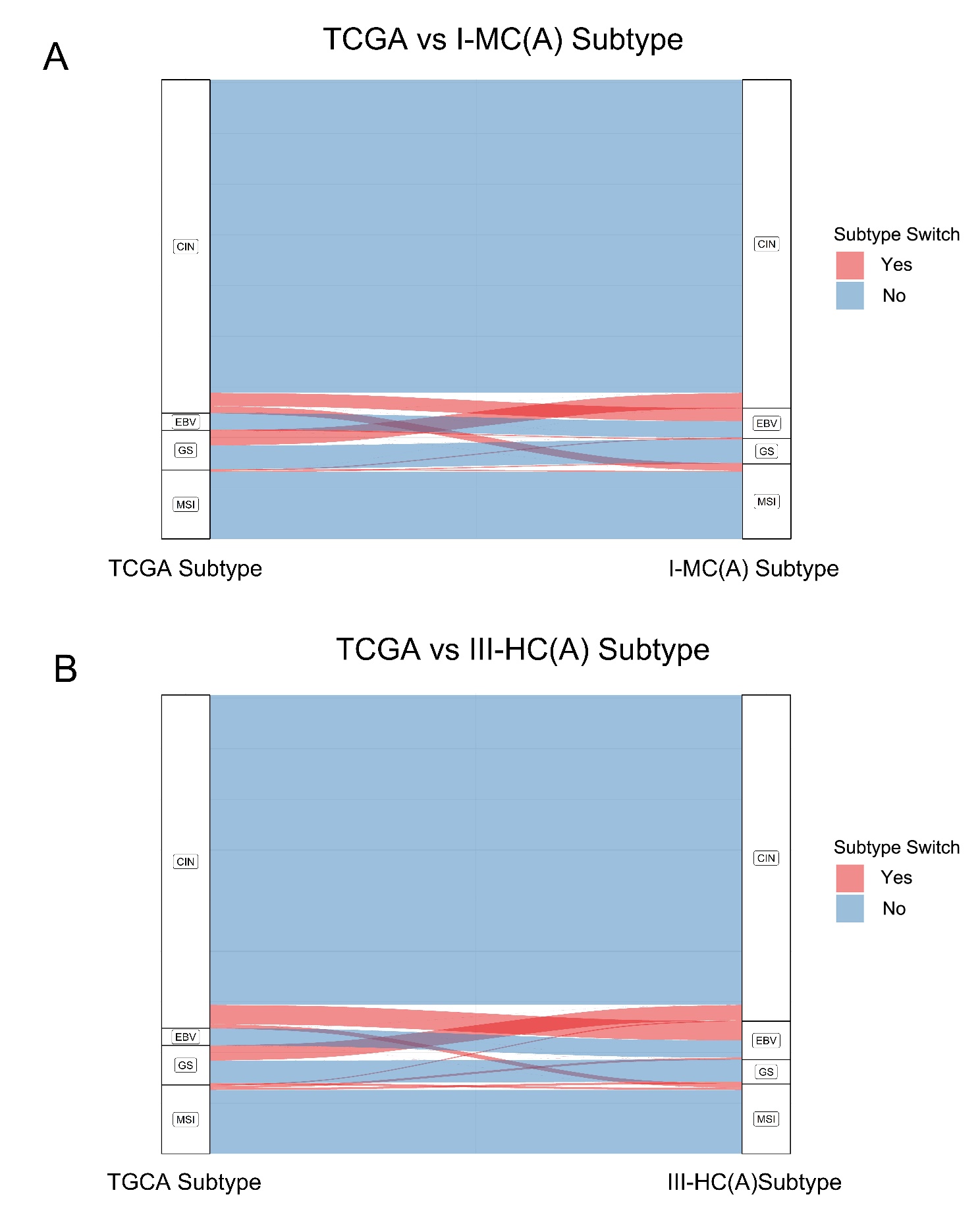
**Supplementary Figure 1. Alluvial plots. (A)** Alluvial plot between TCGA subtypes and I-MC(A) subtypes. **(B)** Alluvial plot between TCGA subtypes and III-HC(A) subtypes.

**2. Supplementary Tables**

**Supplementary Table. 1** Multivariate survival analyses using all samples from TCGA and I-MC(A), II-HC(A) and III-HC(A) defined subtypes (Age and Sex).

| **Classifier** | **Factors** | **Characteristics** | **Count (%)** | **95% CI** | **Hazard Ratio** | **P-value^1^** |
| --- | --- | --- | --- | --- | --- | --- |
| TCGA | Age | <65 (Reference) | 188 (43.9%) | / | / | / |
|  |  | >=65 | 240 (56.1%) | (1.02, 1.93) | 1.39 | 0.036* |
|  | Sex | Female (Reference) | 126 (29.4%) | / | / | / |
|  |  | Male | 302 (70.6%) | (0.88, 1.80) | 1.25 | 0.218 |
| I-MC(A) | Age | <65 (Reference) | 188 (43.9%) | / | / | / |
|  |  | >=65 | 240 (56.1%) | (1.07, 2.00) | 1.45 | 0.017* |
|  | Sex | Female (Reference) | 126 (29.4%) | / | / | / |
|  |  | Male | 302 (70.6%) | (0.90, 1.80) | 1.27 | 0.171 |
| II-HC(A) | Age | <65 (Reference) | 188 (43.9%) | / | / | / |
|  |  | >=65 | 240 (56.1%) | (1.06, 2.0) | 1.44 | 0.021* |
|  | Sex | Female (Reference) | 126 (29.4%) | / | / | / |
|  |  | Male | 302 (70.6%) | (0.90, 1.80) | 1.28 | 0.168 |
| III-HC(A) | Age | <65 (Reference) | 188 (43.9%) | / | / | / |
|  |  | >=65 | 240 (56.1%) | (1.07, 2.04) | 1.46 | 0.017* |
|  | Sex | Female (Reference) | 126 (29.4%) | / | / | / |
|  |  | Male | 302 (70.6%) | (0.90, 1.80) | 1.28 | 0.168 |

^1^Statistical significance is based on the fitted multivariate cox model (log-rank test), * if P<0.05.
